# ProtSpace: Protein Universe in Your Browser

**DOI:** 10.64898/2026.05.04.722720

**Authors:** Tobias Senoner, Peyman Vahidi, Tobias Olenyi, Florin Senoner, Gökhan Sisman, Elias Kahl, Burkhard Rost, Ivan Koludarov

**Affiliations:** School of Computation, Information, and Technology (CIT), Department of Informatics, Bioinformatics & Computational Biology, TUM (Technical University of Munich), 85748 Garching/Munich, Germany; Helmholtz Institute of Computational Biology, Computational Health Center, Helmholtz Munich, 85764 Neuherberg, Germany; Valtech, 7 Route de Saint-Julien, CH-1227 Carouge, Geneva, Switzerland; TUM School of Life Sciences Weihenstephan (WZW), Alte Akademie 8, Freising, Germany; Institute for Advanced Study (TUM-IAS), Lichtenbergstr. 2a, 85748 Garching/Munich, Germany

**Author notes:** These authors contributed equally (Tobias Senoner, Peyman Vahidi). These authors share senior authorship (Burkhard Rost, Ivan Koludarov). **Website:** https://protspace.app **Access Statement:** This website is free and open to all users, with no login required. **Code Repository:** https://github.com/tsenoner/protspace_web (Apache License 2.0).

**Keywords:** protein language model, embedding visualization, web application, functional annotation, dimensionality reduction, proteome-scale

## Abstract

Protein Language Models (pLMs) generate per-protein embeddings that encode functional, structural, and evolutionary information, yet the relationships captured in these representations remain difficult to explore systematically. ProtSpace (https://protspace.app) is a web application for interactive visualization of pLM embedding spaces, enabling hypothesis generation directly in the browser without installation. Unlike traditional network-based tools that exclusively visualize amino acid sequence similarity, ProtSpace explores embedding spaces, revealing relationships often not captured by traditional comparisons. Users provide protein sequences or pre-computed embeddings through a Google Colab notebook or the Python CLI; the pipeline applies dimensionality reduction, retrieves 38 annotation types spanning UniProt, InterPro, NCBI Taxonomy, TED structural domains, and sequence-based predictors served via Biocentral, and produces a portable binary file for the browser-based viewer. WebGL-accelerated rendering supports interactive exploration of over 570,000 proteins. Distinctive features include per-point pie charts for multi-label annotations and integrated 3D structure viewing through AlphaFold2 predictions. All computation happens on the user’s machine, ensuring data privacy. We demonstrate the utility of ProtSpace through a progressive zoom-in across biological scales: from global proteome organization of Swiss-Prot, through cross-species comparison revealing conserved and lineage-specific families, to functional hypothesis generation within the beta-lactamase superfamily. ProtSpace is freely available at https://protspace.app under the Apache 2.0 license.

**Key points:**

1. ProtSpace is a free, open-source web application that visualizes protein Language Model (pLM) embeddings as interactive maps, scaling to 570,000 proteins entirely client-side.
2. A zero-installation Google Colab notebook and a Python CLI prepare visualization-ready bundles from FASTA files, UniProt queries, or pre-computed HDF5 embeddings, automatically retrieving 38 annotation types from five sources (UniProt, InterPro, NCBI Taxonomy, TED structural domains, and Biocentral sequence predictors) alongside custom CSV metadata.
3. Application examples demonstrate that embedding visualizations generate testable biological hypotheses at multiple scales, from proteome-wide organization through species-level comparison to family-level functional discovery, and that these are complementary to traditional sequence-based analyses.

## Introduction

Protein Language Models (pLMs) such as ProtT5 [1], ESM-2 [2], and Ankh [3] learn rich sequence representations that implicitly capture evolutionary signal, which comes from self-supervised training on hundreds of millions of natural protein sequences. The resulting per-protein embeddings have driven advances in function annotation [4], property prediction [5], and remote homology detection into the midnight zone [6], underscoring the growing role of pLMs across biology. Embedding-based approaches have been applied to visualize microbiome protein spaces [7], estimate sequence conservation at functional sites without alignment [8], and navigate enzyme sequence spaces for biocatalyst discovery [9].

Existing visualization tools each address different aspects of this challenge (Table S1; Supporting Online Material). CLANS [10] and EFI-EST [11, 12] visualize sequence-similarity networks derived from BLAST [13] rather than learned representations, and neither provides integrated, switchable protein annotations: CLANS requires manual group definition, while EFI-EST embeds UniProt attributes accessible only through Cytoscape’s visual mapping interface. CLANS depends on a local Java Runtime Environment, while EFI-EST depends on Cytoscape, recently also available as a browser application [14]. General-purpose embedding viewers such as TensorFlow Embedding Projector [15] and Embedding Atlas [16] run in the browser. Embedding Atlas scales to millions of points with automatic clustering but provides no protein-specific annotations, evidence codes, multi-label support, or structure viewing. PLVis [17] offers pLM projections for reference proteomes with cluster enrichment analysis, but is restricted to pre-computed embeddings and lacks runtime annotation switching, multi-label visualization, evidence codes, and 3D structure viewing. ProtSpace addresses these gaps: it retrieves 38 annotation types with one-click switching from UniProt [18], InterPro [19], NCBI Taxonomy [20], TED structural domains [21], and Biocentral sequence predictors [22], extensible with custom CSV metadata for user-defined expert features. It visualizes multi-label annotations as per-point pie charts, displays ECO evidence codes [23] inline, integrates 3D structure viewing via Mol* [24], and explores over 570K proteins entirely in the browser, all from raw sequences without requiring local software.

Our original ProtSpace Python package [25] made pLM embedding spaces accessible to experimental biologists, enabling them to map unknown proteins (e.g., from novel transcriptomes or proteomes) against known families and generate functional hypotheses visually [26, 27]. However, community feedback identified three limitations that motivated a complete web-native reimplementation: installation complexity restricted adoption, performance degraded beyond approximately 10K proteins, and annotations required manual preparation.

Here, we present ProtSpace, a web application that addresses all three limitations (Figure 1). It runs entirely in the browser, eliminating installation requirements. WebGL-based rendering scales to over 570K proteins with sub-second interaction latency (Figure S1). Automatic retrieval of curated and predicted annotations (detailed in Annotation System) eliminates manual data preparation. To our knowledge, ProtSpace is the first tool to combine interactive pLM embedding exploration with integrated functional annotations at cross-proteome scale, entirely in the browser.

**Figure 1:**
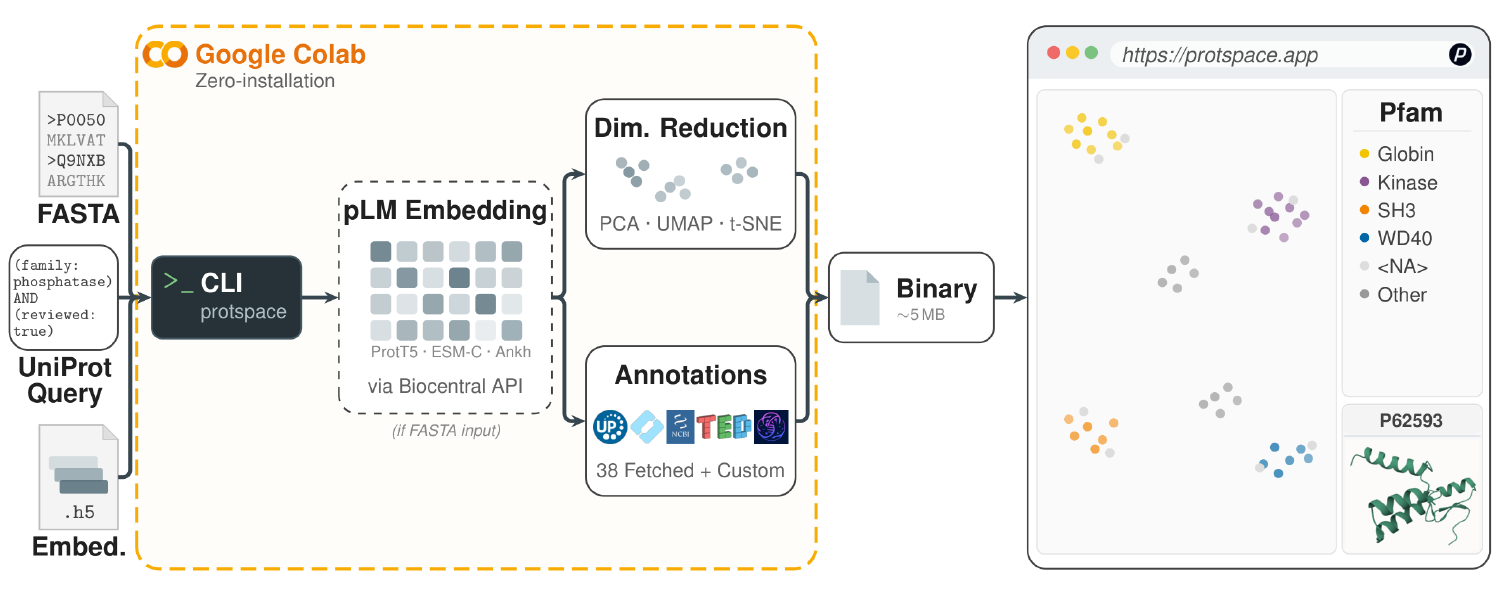
ProtSpace converts protein data into interactive embedding maps. Users provide one of three input types (a FASTA file, a UniProt query, or a pre-computed HDF5 embedding file) to a Google Colab notebook or the Python CLI (`pip install protspace`). For sequence input (provided directly as FASTA or fetched via a UniProt query), the pipeline computes pLM embeddings via the Biocentral API [22] (twelve models available, including ProtT5, ESM-2, Ankh, and ESM-C); pre-computed HDF5 inputs skip this step (dashed box). The pipeline then applies dimensionality reduction (PCA, UMAP, t-SNE, PaCMAP, LocalMAP, or MDS) and retrieves annotations from five public protein data sources, with optional custom CSV metadata for user-defined expert features. The output `.parquetbundle` file (∼5 MB on average) is loaded at https://protspace.app, where the client-side interface provides interactive visualization with annotation coloring, an interactive legend, and integrated 3D structure viewing, without server upload.

### Implementation

#### Architecture and Data Preparation

ProtSpace is implemented as a modular web application using Lit [28] web components. The scatterplot component renders hundreds of thousands of data points via a WebGL-accelerated canvas, maintaining interactive frame rates (Figure S1) for pan, zoom, and selection. A quadtree spatial index enables sub-millisecond point queries for hover and click interactions at maximum scale. All computation and rendering occur client-side: no protein data is uploaded to any server, ensuring complete data privacy.

Data is exchanged in the .`parquetbundle` format (Note S1), which concatenates multiple Apache Parquet [29] tables into a single file containing annotations, projection coordinates, metadata, and optional visualization settings. This binary format achieves high compression ratios (the entire Swiss-Prot dataset compresses to ∼45 MB - for 2 dimensionality reduction methods + 38 annotation types) while supporting efficient partial reads.

Two data preparation paths serve users with different levels of computational expertise (Figure 1). Both accept three input types: FASTA files and UniProt queries are embedded via the Biocentral API [22] (twelve pLMs available: ProtT5, ProstT5, five ESM-2 variants, three Ankh models, and two ESM-C models), while pre-computed HDF5 embeddings from any pLM are used as-is. The Google Colab notebook provides a guided interface: users choose dimensionality reduction methods and annotation types, and the notebook exports a `.parquetbundle` file. UniProt itself provides pre-computed ProtT5 embeddings for all entries [18], eliminating the need for GPU resources when working with known proteins.

The **Python CLI** (`pip install protspace`) provides full programmatic control via the `protspace prepare` command. Each pipeline stage is also available as an individual sub-command (`embed`, `project`, `annotate`, and `bundle`) for custom workflows. All intermediate artifacts (FASTA, embeddings, annotations, similarity matrices, and projections) are cached under {`output`}`/tmp/` so that retrying a run only recomputes the stages explicitly invalidated via `--refetch`, and each run writes a YAML `run.log` capturing parameters, protein counts, and stage-level timings for reproducibility. Both paths allow mixing database annotations with custom CSV metadata files.

Six dimensionality reduction methods are available (Table S3): PCA, UMAP [30], t-SNE [31], PaCMAP [32], LocalMAP [33], and MDS [34], each configurable, such as distance metric and random seed. Unlike the initial ProtSpace Python package [25], which supported both 2D and 3D projections, the web application produces 2D projections exclusively. Despite preserving more variance, 3D projections proved counterproductive in practice. WebGL rendering capacity drops sharply at the point counts we target, and users must constantly rotate the view on a 2D screen, which obscures spatial relationships rather than revealing them. Restricting to 2D improved both visual clarity and interactive performance at proteome scale. Multiple projections are computed simultaneously and can be switched interactively in the viewer without recomputation.

#### Annotation System

ProtSpace retrieves 38 annotation types from five data sources via public APIs (Table S2): 14 from UniProt [18], including EC numbers with enzyme name resolution, Gene Ontology [35] terms split by ontology aspect, subcellular localization, protein families, and sequence length; 10 from InterPro [19], providing domain annotations from Pfam [36], SUPERFAMILY, CATH-Gene3D, SMART, CDD, PANTHER, PROSITE, PRINTS, Phobius signal peptide predictions (each with per-domain bit scores), and a derived Pfam-to-clan mapping; 9 taxonomic ranks from NCBI Taxonomy [20], resolved via the UniProt Taxonomy API from UniProt organism identifiers; 1 structure-based annotation (`ted_domains`) from TED, The Encyclopedia of Domains [21], via the AlphaFold Database API [37], assigning CATH classifications and pLDDT confidence to predicted structural domains; and 4 sequence-based predictions served by the Biocentral API [22], specifically subcellular localization, membrane/soluble status, signal peptide, and transmembrane topology, enabling annotation of proteins absent from curated databases. Evidence codes from the Evidence and Conclusion Ontology (ECO) [23] are preserved for six UniProt-sourced annotation fields (subcellular location, EC number, the three GO aspects, and protein family); hovering over a protein displays its evidence codes (e.g., EXP for experimental, IDA for direct assay) alongside annotation values in the detail tooltip, allowing users to distinguish experimentally verified from computationally inferred annotations. Beyond these public sources, users can always supply their own expert annotations as custom CSV metadata, which are merged with retrieved values and take precedence on name collisions, supporting proprietary experimental data or user-defined category schemes.

#### Interactive Visualization

Users load `.parquetbundle` files by drag-and-drop or via Import button. The viewer supports annotation coloring for both single-label and multi-label annotations. A distinctive feature is the display of multi-label annotations as per-point pie charts: for example, a protein annotated with three Pfam domains appears as a three-slice pie at its embedding position, revealing domain architecture patterns across entire datasets.

The legend system supports drag-and-drop reordering, per-category visibility toggling, shape and color assignment, and selection from six color palettes, defaulting to Kelly’s maximum-contrast scheme [38] with colorblind-safe alternatives including Okabe-Ito [39].

All settings persist via browser storage and can be exported within the `.parquetbundle` for reproducible sharing.

Integrated 3D structure viewing uses Mol* [24] to auto-load AlphaFold [40] predicted structures, retrieved through the 3D-Beacons API [37, 41], with direct links to the AlphaFold Database, UniProt, and InterPro entries. Additional features include protein ID search with multi-ID queries, isolation and filter modes for subgroup analysis, per-category shape assignment, and export in PNG, PDF, protein ID list, or `.parquetbundle` formats with customizable dimensions. Exported `.parquetbundle` files preserve the user’s legend configuration (colors, shapes, sort order, and visibility), enabling exact reproduction and sharing of curated views with collaborators.

#### Performance

To evaluate client-side responsiveness across diverse hardware, we developed an automated WebGL benchmark suite that measures rendering performance for four interaction scenarios (annotation switching, zoom, pan, and point selection) across nine datasets ranging from 800 to 248K proteins (Figure S1). We collected results from 12 participants using 3 browser engines (Chromium, Firefox, Safari) on machines with different GPUs and CPU architectures. Pan and zoom render in ∼1 ms regardless of dataset size, confirming that camera transformations are independent of point count. Annotation switching and point selection scale linearly: median render times remain below 10 ms at 5K proteins, below 145 ms at 152K, and below 230 ms at 248K, well within interactive thresholds.

### Application Examples

We illustrate ProtSpace’s capabilities through a progressive zoom-in across biological scales: from a global view of all reviewed proteins, through a two-species comparison, to a detailed analysis of a single protein superfamily.

#### Global Proteome Organization

To demonstrate scalability, we visualized all 573K Swiss-Prot proteins colored by domain of life using UMAP projection (Figure 2A). UMAP resolves this space into largely distinct bacterial and eukaryotic clusters with limited overlap, while archaeal and viral proteins occupy compact regions. The restricted viral footprint likely reflects smaller proteome sizes, and the compact archaeal representation may stem from their underrepresentation in pLM training datasets, highlighting a potential direction for targeted sequencing efforts.

**Figure 2:**
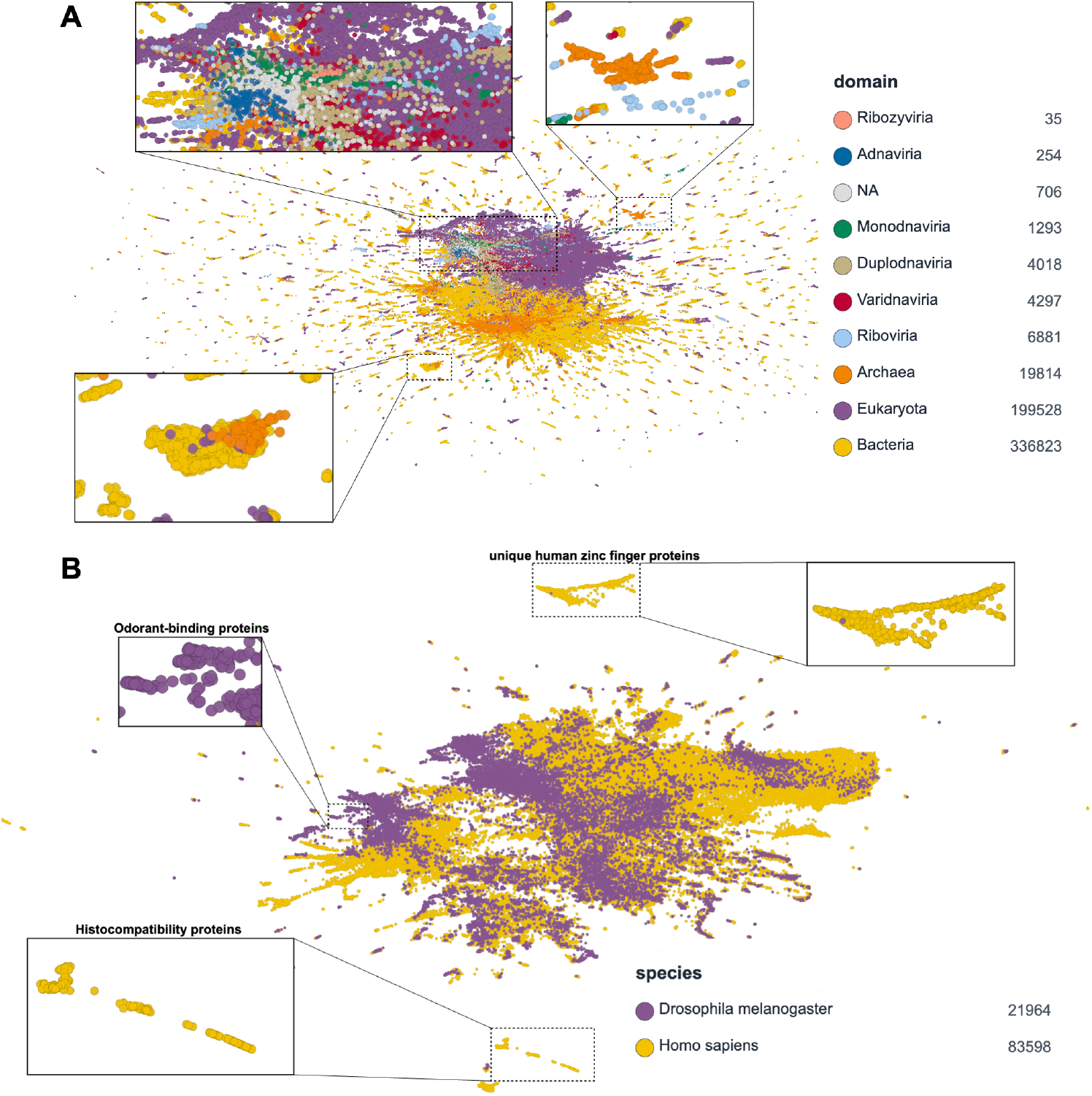
From global proteome organization to species-level comparison. Pre-computed ProtT5 [1] embeddings were retrieved from UniProt; UMAP [30] projection and annotation retrieval were performed with the protspace Python CLI. (A) 573K Swiss-Prot [18] proteins colored by domain of life: distinct bacterial and eukaryotic clusters emerge; archaeal and viral proteins occupy compact regions. (B) Joint projection of the *Homo sapiens* and *Drosophila melanogaster* reference proteomes (105K proteins) colored by species reveals conserved protein families in overlapping regions and species-specific families in isolated clusters.

#### Cross-Species Proteome Comparison

Zooming from the global view into two species, a joint projection of the *Homo sapiens* and *Drosophila melanogaster* reference proteomes (105K proteins) shows extensive overlap alongside regions dominated by a single species (Figure 2B). Visual clusters alone are not evidence of biological signal (random vectors also form clusters under UMAP), so ProtSpace treats this view as a starting point for interactive hypothesis generation rather than as a stand-alone claim. Recoloring the same projection at https://protspace.app by protein family shows that conserved families occupy the overlap region (e.g., 1,703 protein kinases across the joint proteome), while species-specific families concentrate in the isolated regions: MHC classes I and II, beta-defensins, and CC chemokines for human; odorant-binding proteins (PBP/GOBP family) and Yellow-family proteins for *Drosophila*.

#### Functional Family Discovery in the Beta-Lactamase Superfamily

Zooming further into a single protein family shared across distant taxa, we examined 152K beta-lactamase superfamily proteins using ProtT5 embeddings with UMAP projection (Figure 3A). Coloring by protein family shows that clusters correspond directly to known beta-lactamase families, including Ambler classes A through D [42], indicating that the embedding captures functionally meaningful features. Because ProtSpace allows users to switch interactively between annotation layers, the same projection can be recolored, for example by EC number or taxonomy, to distinguish functional from phylogenetic signal without recomputation.

**Figure 3:**
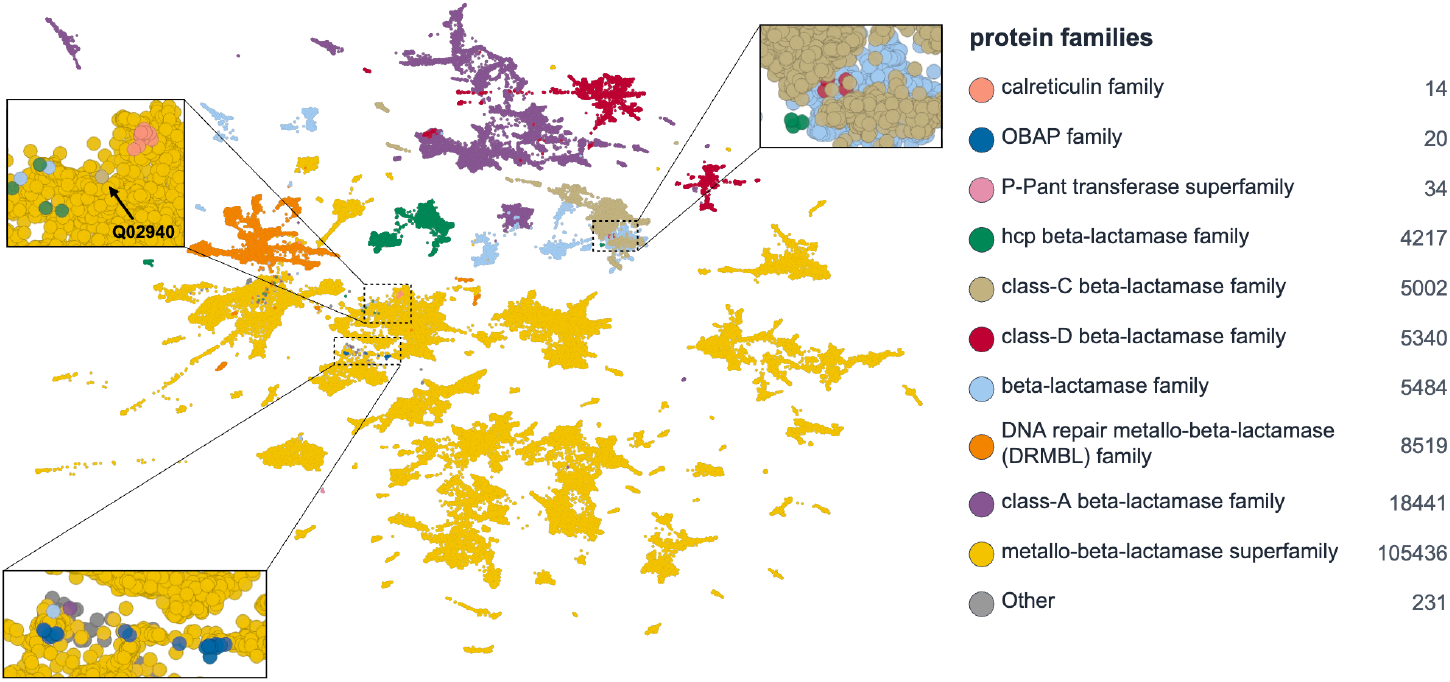
Functional families emerge in beta-lactamase embedding space. Proteins were retrieved from UniProt using the query `family:`”`beta-lactamase`” (152K entries); ProtT5 embeddings were obtained from UniProt, UMAP projection applied, and annotations fetched via the protspace Python CLI. (A) Colored by protein family: clusters correspond to known families (Ambler classes A–D); proteins with generic annotations co-cluster with defined families; annotated proteins outside expected clusters flag candidate misannotations; and clusters without defined family correspondence suggest novel families. (B) Same projection colored by kingdom taxa; few clusters are restricted to a single domain of life, indicating functional rather than phylogenetic clustering.

Several observations generate testable hypotheses. First, proteins that are either unannotated or carry only generic labels (e.g., “Uncharacterized protein”) co-cluster tightly with well-characterized families, enabling putative reassignment based on embedding neighborhood. Second, approximately 860 proteins (∼0.7% of the dataset) with specific family annotations fall outside their expected clusters (fewer than half of their 50 nearest UMAP neighbors share the same annotation), flagging candidate misannotations or atypical evolutionary histories. For example, Q02940, annotated as a class-C (serine) beta-lactamase, clusters instead with the metallo (Zn^2+^) beta-lactamases in both UMAP and PCA projections, a candidate for reassignment across mechanistic classes. Third, multiple compact and well-separated clusters lack correspondence to any currently defined family, representing candidates for novel family descriptions.

Coloring by kingdom taxa (Figure 3B) confirms that phylogenetic signal contributes minimally to the clustering: few clusters contain proteins from only a single domain of life, indicating that the embedding captures functional rather than phylogenetic relationships.

## Conclusion

ProtSpace provides accessible, scalable, and annotation-rich exploration of pLM embedding spaces through a fully browser-based interface. To our knowledge, it is the only web application combining interactive pLM embedding visualization with integrated annotations from five complementary sources (spanning curated sequence databases, predicted structural domains, and learned sequence-based predictors) at proteome scale. The application examples, from global proteome organization through cross-species comparison to functional family discovery, illustrate the tool’s capacity to generate testable biological hypotheses across multiple scales. Current limitations point directly to future work: scaling beyond hundreds of thousands of proteins, where browser memory and WebGL buffers become bottlenecks; the lack of incremental projection, since most dimensionality reduction methods alter the global layout when new proteins are added, requiring full recomputation; and adding built-in cluster detection with enrichment statistics and density contours to complement the current visual exploration. Integration of newer pLM architectures beyond the twelve currently served through Biocentral will keep ProtSpace aligned with the rapidly evolving model landscape. ProtSpace is freely available at https://protspace.app. All figures for this publication were created in-app.

## Abbreviations

DR: dimensionality reduction
EC: Enzyme Commission
ECO: Evidence and Conclusion Ontology
GO: Gene Ontology
PCA: Principal Component Analysis
pLM: protein language model
t-SNE: t-distributed Stochastic Neighbor Embedding
UMAP: Uniform Manifold Approximation and Projection

## Data Availability

ProtSpace is freely available at https://protspace.app. Source code is available at https://github.com/tsenoner/protspace_web (Apache 2.0). The Python data preparation package is installable via `pip install protspace`. All `.parquetbundle` files used to generate Figures 2–3 are available for download at [**TODO: insert persistent URL, e.g**., **Zenodo DOI**]. Example datasets and the Google Colab notebook are linked from the website.

## Acknowledgements

We thank Maria Martin and Alex Bateman as well as Matthias Blum, Aurelien Luciani, Abdulrahman Hussein, and Yvonne Lussi (EMBL-EBI) for constructive discussions and feedback. We thank Nikita Kugut (TUM) for his support with many aspects of this work. We also thank the volunteers who ran performance benchmarks on their machines. We thank the Technical University of Munich (TUM) for providing facilities and resources. Finally, we thank those who deposit experimental data in public databases, maintain these databases, and develop methods to enrich experimental data.

## Supplementary Materials

**Table S1:** ProtSpace is the only tool combining browser-based pLM embedding visualization with integrated protein annotations at proteome scale. Feature comparison across five visualization tools. ProtSpace uniquely offers 38 annotation types from five sources (UniProt, InterPro, NCBI Taxonomy, TED structural domains, and Biocentral predictors), multi-label pie charts, integrated 3D structure viewing, and evidence code display, all running client-side without installation. ^a^Web portal for pre-computed reference proteomes; custom analyses require Google Colab. ^b^Web portal generates sequence similarity networks; visualization requires Cytoscape desktop or Cytoscape Web [14].

| Feature | ProtSpace Web | Emb. Atlas | PLVis | CLANS | EFI-EST |
| --- | --- | --- | --- | --- | --- |
| **Access** | Web browser | Web browser | Web + Colab<sup>a</sup> | Java desktop | Web + Cytoscape<sup>b</sup> |
| **Installation** | None | None | None | Java + BLAST | Cytoscape<sup>b</sup> |
| **Max proteins** | 573,000 | Millions | Full proteomes | ~10,000 | 75,000 |
| **Privacy** | Client-side | Client-side | Server-side | Local | Server-side |
| **Input** | FASTA / Query / H5 | Any embedding | pLM (pre-computed) | BLAST P-values | Sequences / families |
| **DR methods** | 6 | UMAP | 4 | Force-directed | N/A (SSN layout) |
| **Annotations** | 38 types, 5 sources | Auto-clustering | Cluster enrich. | Manual groups | UniProt node attrs. |
| **Custom annot.** | CSV (via CLI) | User metadata | No | Manual | No |
| **Multi-label** | Pie charts | No | No | No | No |
| **3D struct.** | AlphaFold (Mol*) | No | No | No | No |
| **Confidence** | ECO / bit / pLDDT | No | No | No | No |
| **Export** | PNG/PDF/IDs/Bundle | CSV/Parquet | CSV (via Colab) | PNG/EPS/clans | XGMML/TSV |
| **License** | Apache 2.0 | MIT | Not specified | GPL | GPL-3.0 |

**Table S2:**
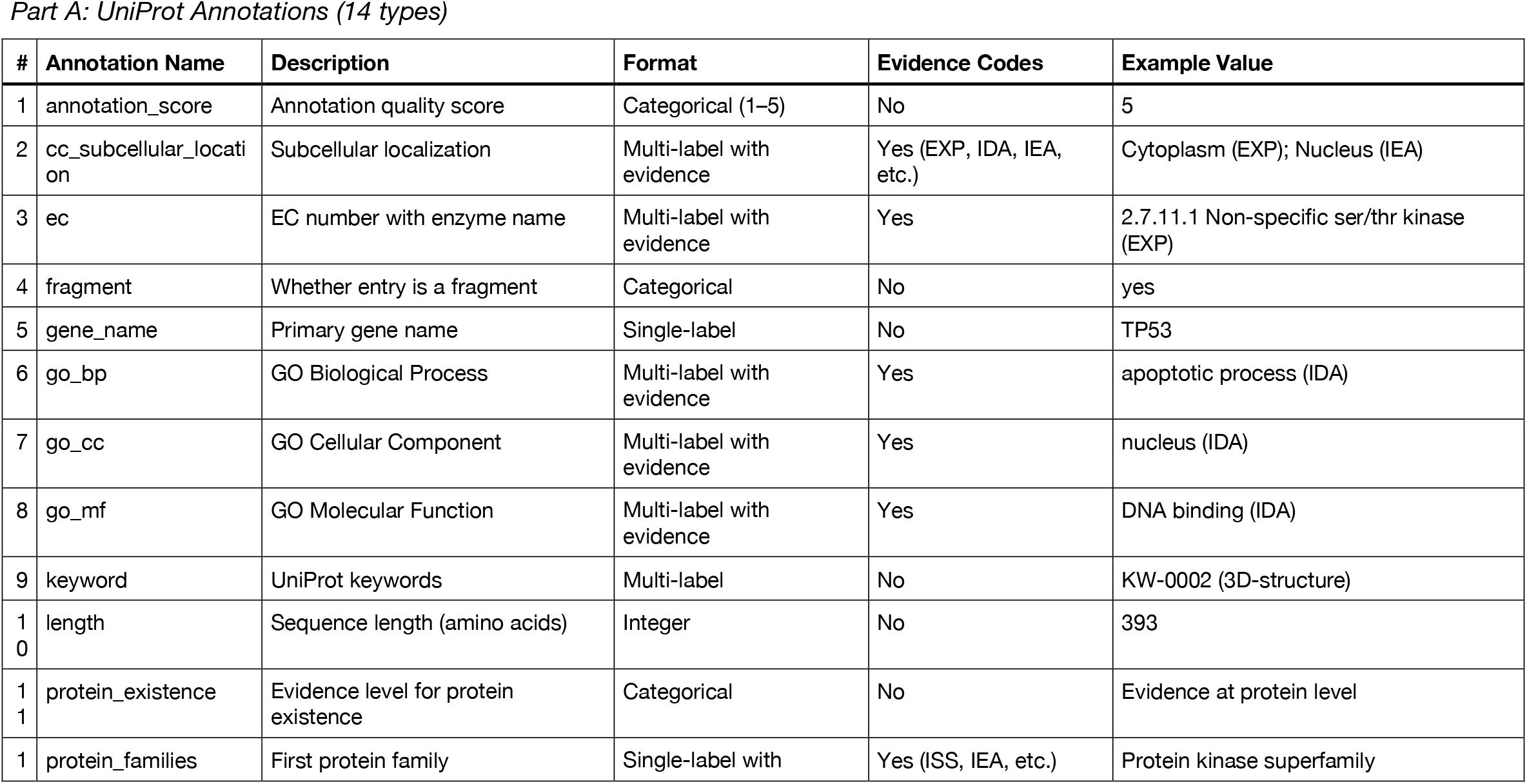
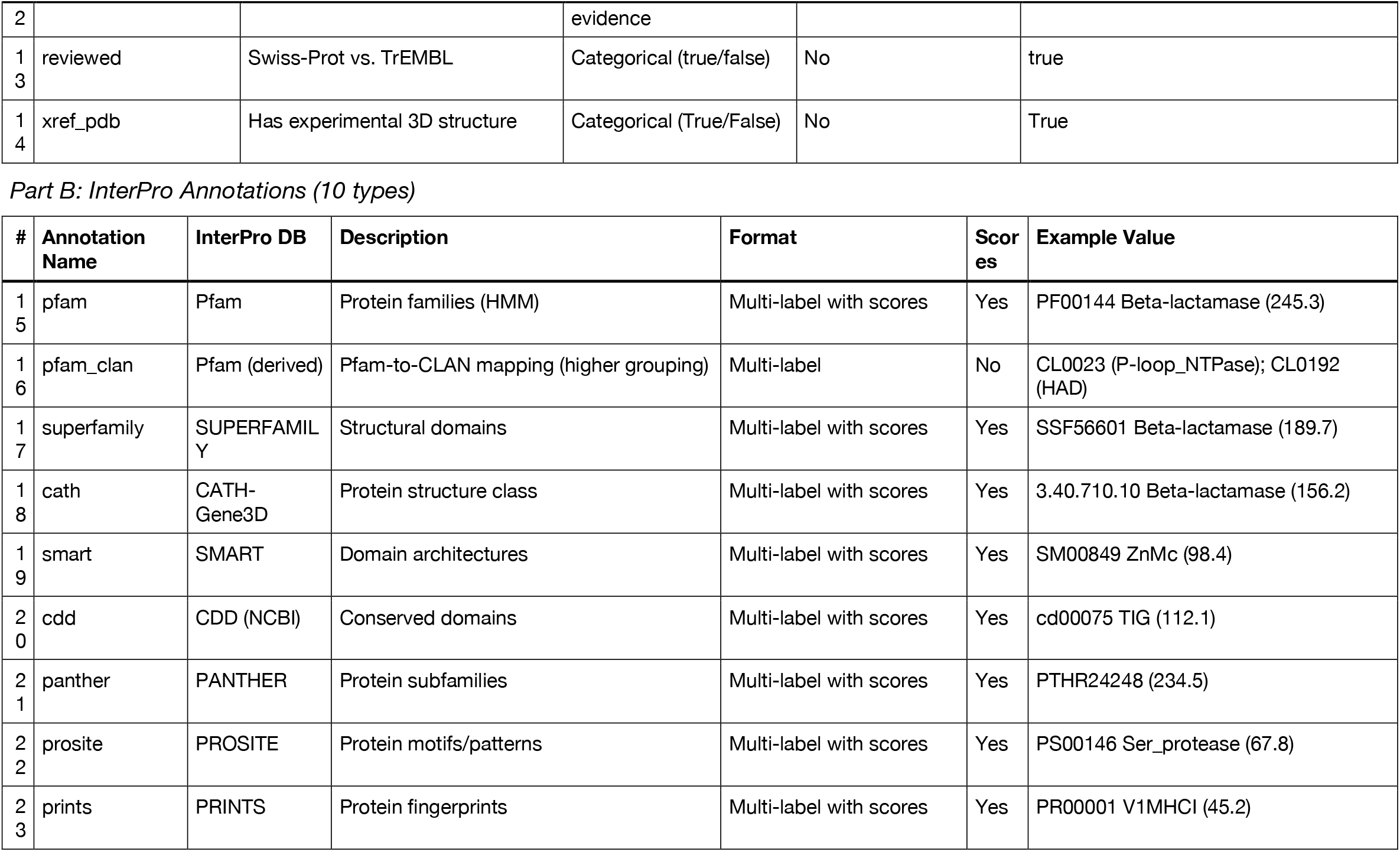
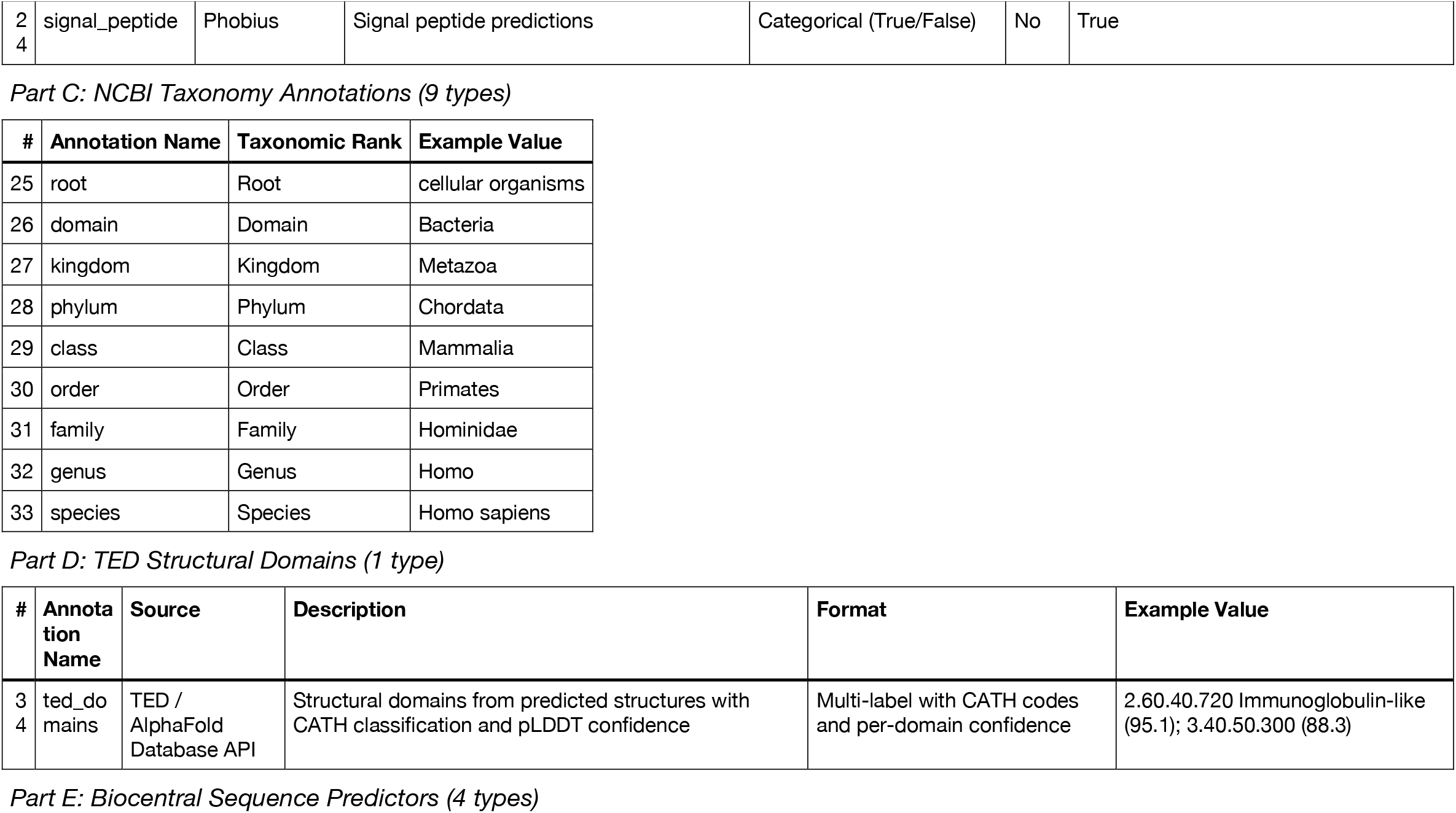
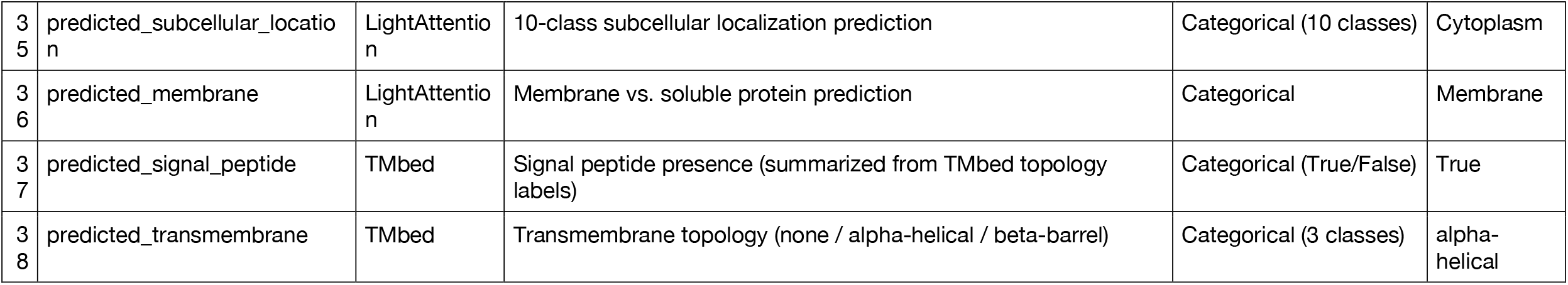
ProtSpace retrieves 38 annotation types from five protein data sources. Annotations are organized by source: UniProt (14 types), InterPro (10 types, including the derived `pfam_clan` mapping), NCBI Taxonomy (9 ranks), TED structural domains via the AlphaFold Database API (1 type), and Biocentral sequence predictors (4 types). Evidence codes from the Evidence and Conclusion Ontology (ECO) are preserved where available for six UniProt-sourced fields. Two additional fields (`protein_name`, `uniprot_kb_id`) are retrieved for tooltip display but are not counted among the 38 selectable types; `gene_name`, formerly tooltip-only, is now a selectable UniProt annotation. Taxonomy is resolved from UniProt organism IDs via the UniProt Taxonomy API (the `taxopy` dependency and associated NCBI database download were removed in v4.1.0); missing ranks are filled with.

**Table S3:** Six dimensionality reduction methods offer complementary views of embedding space. Six methods project high-dimensional pLM embeddings to 2D. All use `random_state = 42` by default for reproducibility; parameters are fully configurable via the Python CLI and Colab notebook.

| Method | Key Parameters | Defaults | Best For |
| --- | --- | --- | --- |
| **PCA** | `n_components` | 2 | Global variance, fast computation |
| **UMAP** | `n_neighbors`, `min_dist`, `metric` | 15, 0.1, euclidean | Balanced local/global structure |
| **t-SNE** | `perplexity`, `learning_rate`, `metric` | 30, 200, euclidean | Cluster-focused visualization |
| **PaCMAP** | `n_neighbors`, `mn_ratio`, `fp_ratio` | 15, 0.5, 2.0 | Hybrid local + global + mid-range |
| **LocalMAP** | `n_neighbors`, `mn_ratio`, `fp_ratio` | 15, 0.5, 2.0 | Local-focused manifold structure |
| **MDS** | `n_init`, `max_iter`, `eps` | 4, 300, 1e-3 | Dissimilarity/distance matrices |

**Figure S1:**
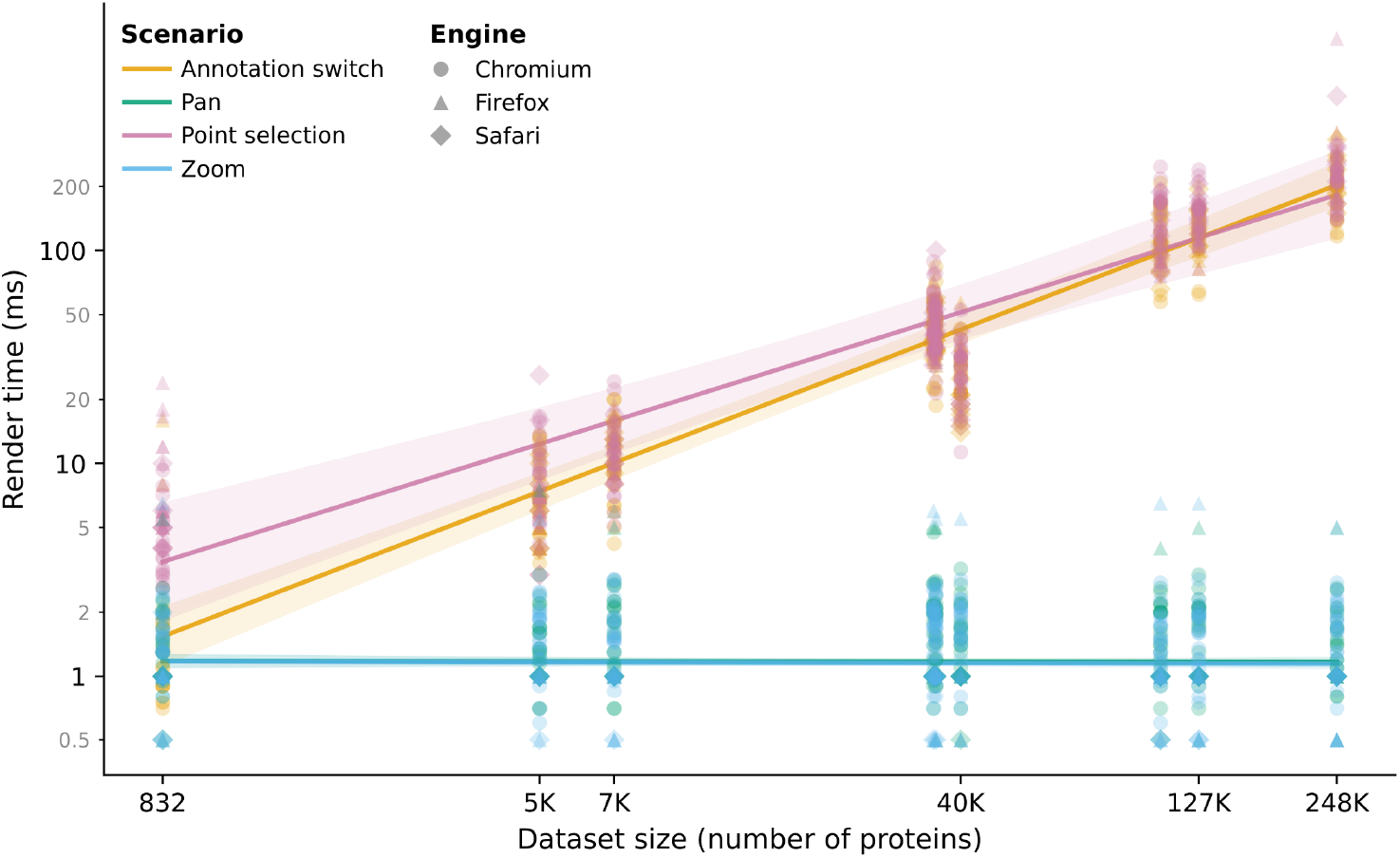
Render time scales linearly with dataset size regardless of browser engine. WebGL render time per interaction scenario (annotation switch, pan, point selection, zoom) across nine datasets (800–248K proteins), measured on 12 participants using 3 browser engines on varied hardware. Browsers are grouped by rendering engine: Chromium (Brave, Chrome, Chromium, Edge, Opera, Vivaldi), Firefox, and Safari. Each point represents the median of three iterations for one dataset–scenario–engine–participant combination; lines show log-linear regression fits with ±2 SE ribbons. Colors follow the Wong colorblind-safe palette [43]. Pan and zoom remain constant (∼1 ms) at all scales; annotation switching and point selection scale linearly with protein count.

### Note @: ParquetBundle Format Specification

A. `parquetbundle` file concatenates 3–4 Apache Parquet tables separated by `---PARQUET_DELIMITER---`:

1. **selected_annotations**: protein identifiers, tooltip fields, and all selected annotation columns
2. **projections_metadata**: projection names, dimensions, and parameters
3. **projections_data**: x/y(/z) coordinates per protein per projection
4. **settings** (optional): a single-row Parquet table whose `settings_json` column holds a JSON-encoded string of per-annotation color, shape, sort order, and visibility configurations

Annotation values use pipe-delimited encoding for multi-label entries and for missing values. Full column-level schema documentation is available in the GitHub repository.

